# Clinical feasibility of spatial transcriptomics using discarded tissue from diagnostic breast biopsies

**DOI:** 10.64898/2025.12.30.696765

**Authors:** Jeffrey P. Sheridan, Quynh Sun, Jiaji G. Chen, Yibing Wei, Veronica Jarzabek, Anirudh Jaishankar, Iqra Amin, Lila Sultan, Thea Nalan, Sarah B. Mueller, Eric J. Burks, Lubna Suaiti, Christopher M. Heaphy, Naomi Y. Ko, Ruben Dries

## Abstract

Spatial transcriptomics holds potential to transform cancer diagnostics, yet significant barriers still limit its clinical translation. First, access to primary patient tissue is often restricted by logistical challenges and patient hesitancy. Second, it remains uncertain whether high-quality spatial transcriptomics data can be generated from clinical biopsy sections as these are collected primarily for diagnostic purposes that do not prioritize RNA-quality. We investigated whether discarded tissue slices, generated during standard pathology procedures, could be repurposed for spatial transcriptomics and alleviate concerns about both tissue availability and quality. Here, we established a pipeline to collect and perform spatial *in situ* transcriptomics on discarded biopsy material from a breast cancer patient, and digitized matched pathology images, including hematoxylin and eosin and traditional histochemistry stains from adjacent sections. Our results show that spatial transcriptomics data from discarded tissue are concordant with the original pathology report, while also providing additional insights such as accurate cell type annotation, detailed spatial architecture, and quantification of biological processes relevant to breast cancer progression. Altogether, our approach using discarded pathology tissue sections provides a practical and scalable solution that would maximize the scientific value of existing clinical specimens and enable high-resolution tumor microenvironment mapping.

## Introduction

Primary patient samples remain essential to advancing our understanding of complex diseases such as cancer. Furthermore, studying primary tumor samples from patients at the molecular and cellular level is often critical in identifying and validating markers of disease progression^1,2^. This also has the potential to uncover new targeted therapeutic modalities tailored for each patient, enabling true personalized medicine^1,2^. However, leveraging the intrinsic potential of these primary samples to this end requires easy-to-implement technologies that can extract the available information of each biospecimen and aid in building large-scale patient datasets. Such efforts are required to ensure the broader population is represented within these samples, whilst accounting for both genetic variation as well as the impact of socioeconomic status on patient outcomes that is required to translate new discoveries into equitable and generalizable patient care.

Emerging spatial transcriptomics technologies have already demonstrated their immense power in the creation of high-resolution biological maps of the tumor microenvironment (TME), allowing researchers and clinicians to map gene expression *in situ* whilst preserving tissue architecture^3,4^. More recent advances have shown a dramatic increase in the robustness of these assays, including generating high-quality data from very small and archived samples, that were previously seen as highly challenging^5^. Furthermore, spatial transcriptomics data can be combined with traditional histopathological evaluation such as hematoxylin and eosin (H&E) or immunohistochemistry (IHC) staining from adjacent sections^6^. These multimodal datasets, including valuable clinical annotations, provide an ideal foundation for the creation of the next generation of digital pathology tools^4^. These new artificial intelligence (AI)-driven methods are poised to maximize the obtained spatial information and improve diagnostic and prognostics capabilities^7^.

Despite this promise, several barriers currently limit the integration of spatial transcriptomics into routine clinical cancer care. Clinical biopsies are processed through time-sensitive, diagnostic workflows that prioritize clinical outcomes, leaving little flexibility for additional research-oriented steps. Moreover, standard clinical processing protocols may compromise RNA integrity, potentially impeding the generation of high-quality spatial transcriptomic data. Collection of additional research biopsies optimized for spatial transcriptomic purposes face several logistical and practical challenges, including storing fresh-frozen specimens and working in RNase-free environments. These include concerns about sample bias and intra-tumor heterogeneity^8–10^, substantial institutional resource requirements^11^, and patient hesitation towards additional invasive procedures, compounded by logistical barriers such as scheduling conflicts.^8,9^ Furthermore, with increasing use of neoadjuvant treatments in breast cancer, the diagnostic biopsy may be the only option to obtain any treatment naive tissue for research purposes. Identifying solutions that are compatible with established clinical workflows could increase tissue availability for research while advancing our understanding of complex diseases improving clinical care.

In this study, we investigated whether otherwise discarded tissue sections from clinical breast cancer biopsies could be repurposed for high-quality spatial transcriptomics analysis. Leveraging a core needle biopsy and established seqFISH technology, we demonstrate that these routinely discarded sections processed through standard clinical histopathology workflows (i.e., formalin-fixed paraffin-embedded tissue) yield robust spatial transcriptomic data that are concordant with pathology reports while providing enhanced characterization of the tumor microenvironment. Importantly, this approach integrates seamlessly into existing pathology workflows with minimal burden on clinical staff and requires no specialized RNAse-free handling beyond routine laboratory practices. Collectively, our findings establish a practical framework for incorporating spatial transcriptomics into routine clinical practice, which could facilitate the development of spatially-informed diagnostic tools and advance precision oncology that is generalizable to other solid tumor cancers.

## Results

### Study design: Accessing discarded clinical biopsy material to overcome clinical barriers in spatial transcriptomic data generation

To evaluate the clinical feasibility of spatial transcriptomics for breast cancer, we first considered the principal barriers limiting its routine integration into oncology workflows, including patient hesitancy to undergo additional invasive procedures^8^, limited availability of biopsy material for research^9^, and logistical constraints associated with patient recruitment and sample collection (**Fig. 1A & Supplemental Fig. 1**). Spatial transcriptomics methods typically require a single section, not whole biopsies to generate high quality data. To overcome these challenges, we sought to establish a framework that directly repurposes discarded tissue sections from breast cancer tissue biopsies. During standard diagnostic workflows, core biopsies are often serially sectioned at multiple levels, with intervening sections typically discarded once adequate tissue representation is obtained (**Fig. 1B**). We recovered these otherwise-discarded sections that have underwent histological processing and placed them onto specialized slides compatible with sequential fluorescence *in situ* hybridization (seqFISH), a high-plex and established *in situ* spatial transcriptomic assay^12^ (**Fig. 1C, Supplemental Fig. 2A**). Importantly, this approach integrates seamlessly with existing pathology protocols and places minimal additional burden on histotechnologists. Notably, we did not perform any steps to mitigate RNA degradation that is typically performed during sample preparation for spatial transcriptomics. The only added step is transferring discarded tissue sections onto specialized slides that are indistinguishable in shape and handling from the standard slides already in routine use. By utilizing otherwise discarded material, this approach maximizes the information yield from each biopsy, enabling systematic incorporation of spatial transcriptomics into routine pathology. If successful, this approach could promote the development of novel digital pathology methods, facilitate the construction of large-scale and genetically diverse biobanks that would better represent all populations, and could serve as a starting point for future precision oncology and personalized therapeutic decision-making (**Fig. 1D**).

**Figure 1.**
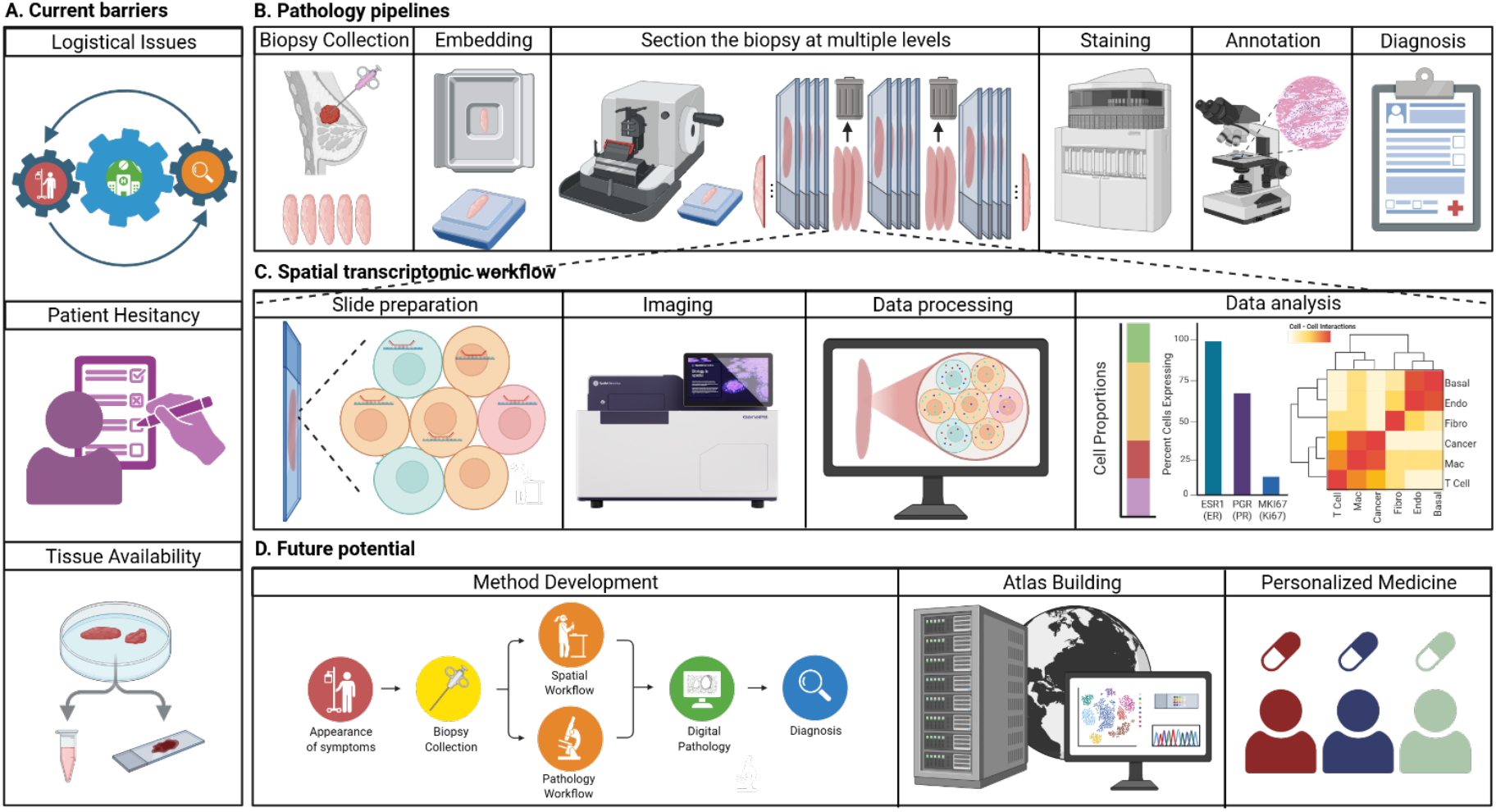
Schematic overview of implementation barriers, approach, and potential for spatial transcriptomics with patient samples. **A**. Current barriers to obtaining primary tissue samples for research. **B**. Clinical procedure for breast cancer biopsy and diagnosis, including sample collection, tissue preparation, generation of discarded tissue sections, immunohistochemistry staining, annotation and diagnosis. **C**. Spatial transcriptomic data generation from discarded tissue sections. **D**. Integration of spatial transcriptomics into diagnostic workflows would enable method development, building atlas-scale datasets, and support spatially informed precision medicine.

### High-quality spatial data generation from discarded tissue slice

To assess data quality from discarded tissue sections, we performed seqFISH with Spatial Genomics, using a 493-gene immune-oncology panel on a breast cancer core needle biopsy from a patient with a ER/PR+, HER2-, KI67-low cancer (**Supplemental Table 1**; **Fig. 2A**). Cell segmentation identified 54,266 cells, comparable to the 58,169 cells detected in an adjacent H&E-stained section. Quality filtering (see methods) retained 97% of cells (n = 52,791) with a median of 84 unique features per cell (**Fig. 2B**). Unsupervised clustering revealed 9 distinct cell populations with characteristic marker expression profiles (**Fig. 2C-D, Supplemental Figs. 3&4**). Luminal cancer cells (74.5%) comprised the majority of the tissue and included a distinct subpopulation (2.7%) of highly proliferative cells marked by high levels of E2F1 and HELLS (**Fig. 2D, Supplemental Fig. 4)**, both known markers of highly proliferative cancer cells^13,14^. The non-cancer fraction within the tumor microenvironment (TME) was comprised of macrophages (6.6%), fibroblasts (4.8%), T cells (4.5%), endothelial cells (2.3%), pericytes (2.0%), mast cells (1.4%), B cells (0.7%), and basal epithelial cells (0.4%). Spatial mapping of these cells revealed tissue organization and structures that are typical of breast cancer biopsies, including ductal carcinoma in situ and milk duct structures (**Fig. 2E**), that matched the routine H&E performed for diagnosis (**Fig. 2F**), demonstrating that discarded diagnostic tissue yields high-quality spatial transcriptomic data representing breast cancer biology.

**Figure 2.**
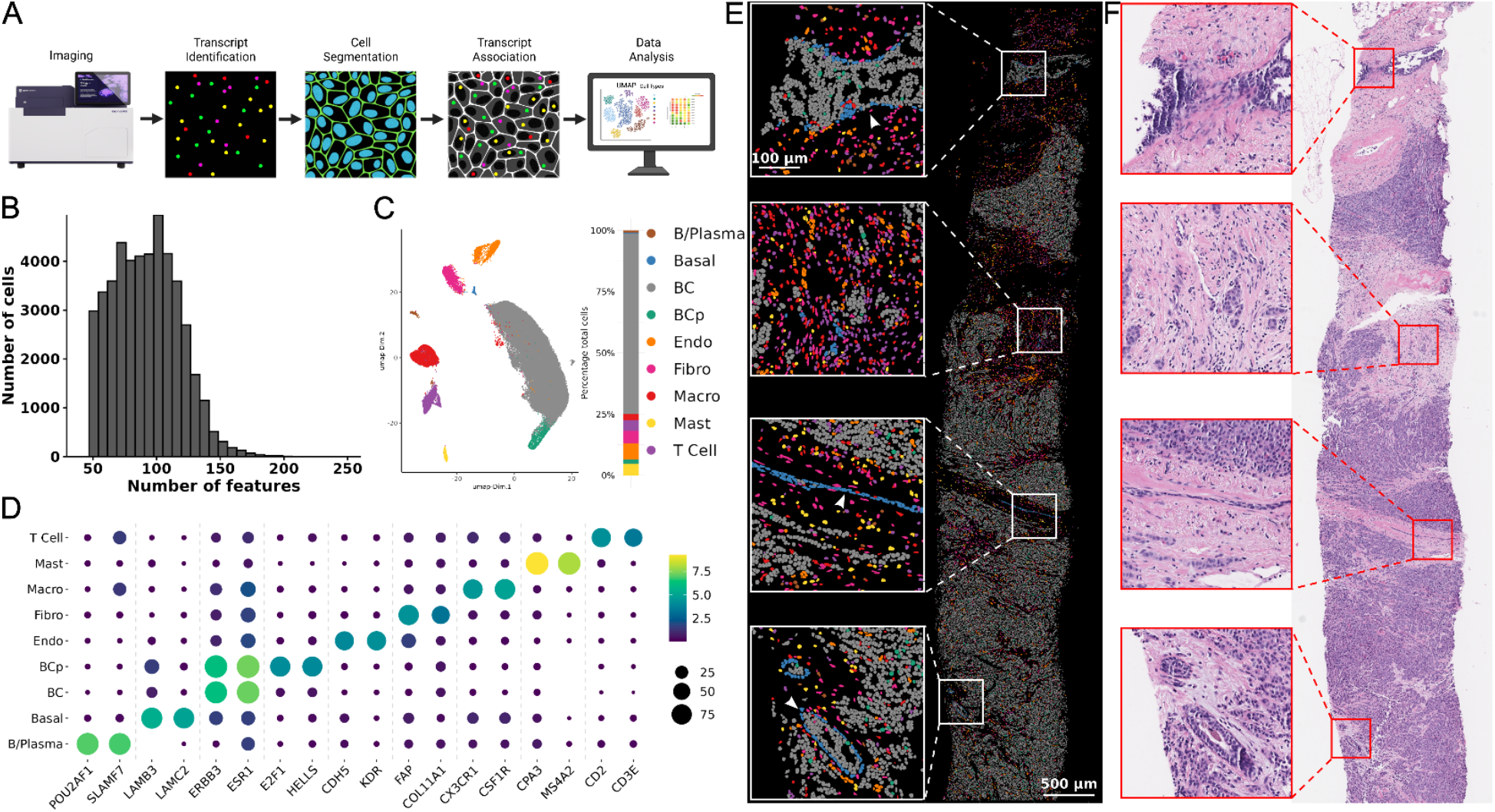
Spatial transcriptomics on discarded tissue biopsy sections. **A**. Overview of the spatial transcriptomics data generation and processing steps. **B**. Histogram displaying the number of unique features (transcripts) detected per cell. **C**. Uniform manifold approximation and projection (UMAP) of cells colored by annotated cell type. Stacked bar plot shows associated cell type proportions within the biopsy. Abbreviations are as follows: B Cells/Plasma Cells (B/Plasma), basal cells (Basal), luminal breast cancer (BC), proliferative luminal breast cancer (BCp), endothelial (Endo), fibroblasts (Fibro), macrophages (Macro), mast cells (Mast). **D**. Dot plot showing representative marker genes for each cell type. **E**. Spatial plots showing the distribution of cell types. **F**. Matched clinical H&E showing matched regions to the spatial map (E).

### Integration of spatial transcriptomics data demonstrates concordance with breast cancer pathology report

To validate if spatial transcriptomic data generated from a discarded breast cancer tissue section can recapitulate clinical diagnostic information, we directly compared spatial gene expression patterns of key breast cancer biomarkers, including ESR1 (estrogen receptor), PGR (progesterone receptor), and MKI67 (Ki67), with immunohistochemistry (IHC) results using the closest relevant tissue sections from the patient’s pathology report. HER2, another common breast cancer marker, was not analyzed since ERBB2, the transcript that encodes HER2, was not present in the gene panel. The diagnostic report indicated ER and PR positivity (>90% for both) with low proliferative index (Ki67 of 15%), findings that were visually confirmed by the associated IHC staining (**Fig. 3A&B**). On the other hand, spatial transcriptomic analysis, performed on a serial section (> 100 µm distance (**Supplemental Fig. 5**)), revealed that 96% and 64% of epithelial cancer cells expressed ESR1 and PGR, respectively, while 7% expressed MKI67 (**Fig. 3A&B**). Hence, spatial expression patterns were concordant with IHC staining across serial sections, with quantitative differences likely reflecting the higher sensitivity of transcript detection compared to protein immunostaining. These results demonstrate that spatial transcriptomics performed on discarded clinical tissue sections provide diagnostic information that is concordant to standard clinical IHC.

**Figure 3.**
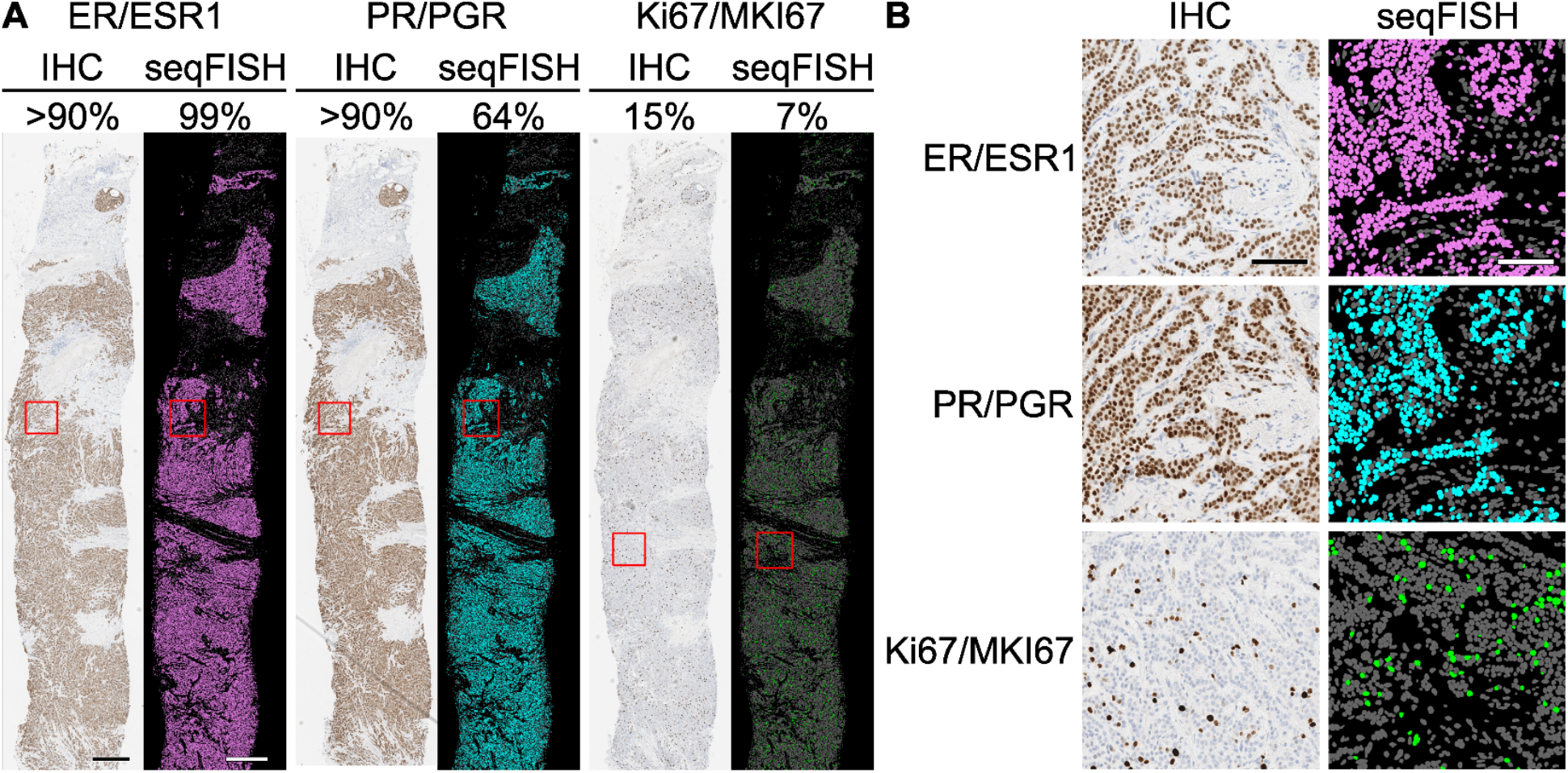
Comparative analysis of IHC and spatial transcriptomics on serial sections. **A**. Shown is expression for IHC staining used for diagnosis (left) and transcripts (right) for closely matched regions on the same biopsy. Percentages on top indicate the positive cells for the relevant marker. IHC was determined during patient diagnosis, whereas seqFISH was calculated by the number of cancer cells expressing the relevant marker divided by the total number of cancer cells. **B**. representative zoomed sections as indicated in A with the relevant red box.

### Tumor niche analysis shows evidence of general immune exclusion

Our results demonstrated that high quality spatial transcriptomics data can be generated from routinely discarded tissue sections, with findings concordant with official pathology reports and histological stains. In addition, the spatial transcriptomics data further enabled detailed cell type identification across the entire tissue section (**Fig. 2E & Supplemental Fig. 3**). To characterize how spatial interactions between various cell types shape the tumor microenvironment, which has been shown to influence clinical outcomes^15–17^, we performed cell-cell interaction analysis on the whole sample (**Supplemental Fig. 6)**. Hierarchical clustering on the enrichments of pairwise cell-to-cell interactions (**Fig. 4A**) showed two main interaction modules. The first module is divided into a cancer fraction that shows preferential proximity between luminal epithelial cancer cells and proliferative luminal epithelial cancer cells, suggesting a relatively homogenous tumor microenvironment dominated by cancer-cancer interactions and the presence of actively proliferating cancer cells at small, localized regions (**Supplemental Fig. 7**). The second module shows spatial interactions between immune cells and fibroblasts, with the strongest interactions observed between the immune cells themselves, including mast cells, T cells, and B cells/plasma cells, which are found predominately in the stromal tissue regions (**Fig. 4A&B**). Together, these results suggest that this is mostly an immune-cold tumor where immune cells are mostly excluded from tumor regions, with T cells, B cells and mast cells, predominantly localizing to the periphery of malignant tissue (**Fig. 4B**).

**Figure 4.**
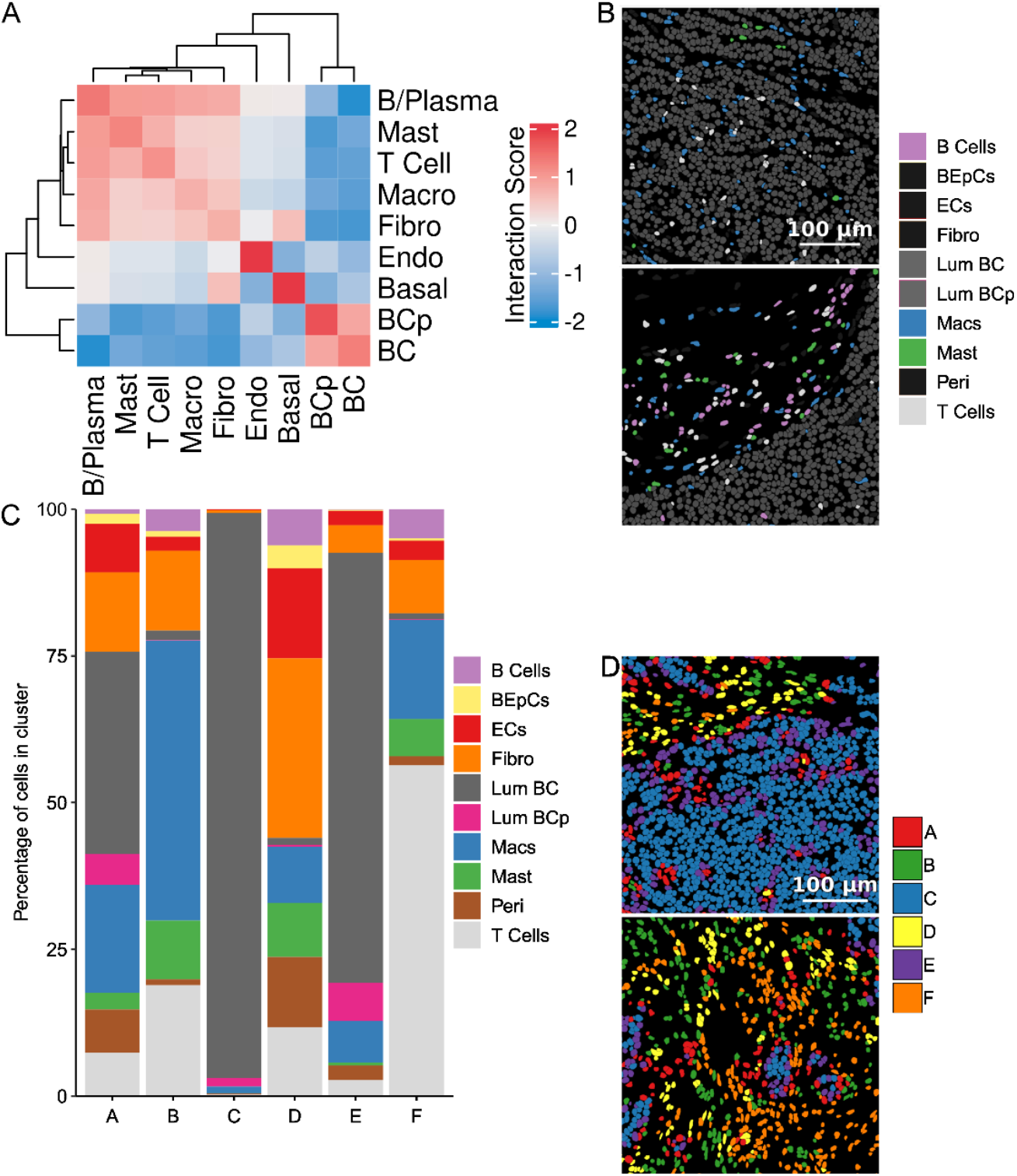
Spatial breast cancer topology analysis at multiple scales. **A**. Heatmap showing interaction scores for pairwise cell-cell interactions for identified cell types. Positive and negative values indicate high and low proximity correlation between cell types, respectively. **B**. Spatial representation of immune cell populations. Cells colored by cell type. **C**. Stacked bar chart showing cell type composition of cellular niche analysis based on cell proximities within the biopsy. **D**. Spatial representation of cellular niches. Cells colored by niche.

The tissue-level analysis characterized broad interaction patterns between pairs of individual cell types. To complement this analysis, we also performed spatial niche analysis, which allows the unbiased identification of more complex and recurrent microenvironmental motifs, consisting of local interactions between multiple cell types. This analysis partitioned cells into six distinct niche clusters (A-F) (**Fig. 4C, Supplemental Fig. 8**). Niche A was the largest (**Supplemental Fig. 9**), comprising approximately 96% luminal breast cancer cells with minimal contributions from other cell types, representing a highly homogeneous tumor-enriched microenvironment (**Fig. 4C**) and in line with our pairwise interaction analysis (**Fig. 4A**). Niche E contained approximately 73% luminal breast cancer cells along with a notable proliferative cancer subset (7%) and macrophages (7%). Niche D showed greater cellular diversity, with luminal breast cancer cells (35%), macrophages (18%), fibroblasts (14%), and endothelial cells (8% each) (**Fig. 4C**). Niche F was characterized by macrophage (30%) and fibroblast (30%) enrichment, along with T cells (15%), and mast cells (10%), representing an immune-infiltrated stromal compartment with minimal cancer cell presence (2%). Niche C displayed a large endothelial population (70%), with small contributions for most other cell types, except for basal and B cells (**Fig. 4C**). Niche B was predominantly T cell-enriched (56%), with additional contributions from macrophages (17%), mast cells (6%), and B cells (5%), and limited cancer cells (1%) (**Fig. 4C**). Spatial mapping revealed that cancer-dominated niches (A, E) occupied the core tumor regions, while immune-enriched niches (B, D, F) localized predominantly to the periphery and stromal areas (**Fig. 4D**). Collectively, these spatial distribution patterns revealed selective immune exclusion from malignant tissue regions. Whereas macrophages showed the broadest distribution across niches, infiltrating both tumor-rich (niche D) and immune-enriched compartments (niches B, D, F), T cells and B cells were largely restricted to peripheral immune niches (B, F), with minimal representation in cancer-dominated regions. These findings suggest active mechanisms within the tumor microenvironment that restrict access of specific immune effector populations capable of suppressing tumor growth and progression^18^.

### Spatial co-expression analysis suggests localized activation of the interferon pathway

Spatial topology analysis showed overall immune exclusion at the cellular level; however, it does not examine how gene expression programs are organized in a spatial and localized manner. Therefore, to capture both transcriptional heterogeneity within cell populations and identify spatially organized expression patterns, we identified spatially variable genes, then using these performed spatial co-expression analysis **(Supplemental Fig. 10)**. This analysis resulted in 7 spatial co-expression modules (**Fig. 5A, Supplemental Fig. 11&12, Supplemental Table 2**). Modules 2 and 3 comprised genes enriched in malignant tissue, while modules 4, 5 and 6, corresponded to mast, T cell, and macrophage signatures respectively. Module 1 was characterized by angiogenesis-related genes localized to the tumor-stroma interface. Interestingly, module 7 exhibited a distinct interferon response signature that was not restricted to a single cell type, despite the overall immune exclusionary environment (**Fig. 5B**). This module included chemokines (CXCL9, CXCL10, CXCL11, CXCL12, CXCL14), interferon-stimulated genes (MX1, PSMB9, TAP2, NLRC5, IDO1), immune cell markers (CD38, SAMF7, NCR1), and immune regulators (IRF4, POU2AF1, NGFR). Interferon responses of this nature have been associated with macrophage and T cell activity and are typically linked to antitumor immunity^19^. Consistent with this interpretation, regions exhibiting module 7 expression contained T cells at greater levels than surrounding tissue, however macrophages were found at a similar level to the surrounding tissue (**Fig. 5C**), which was obscured from our spatial topology analysis (**Fig. 4A&C**). These cells also contributed the greatest to interferon response gene expression pattern (**Fig. 5D**). Diminished interferon signaling has been associated with poor prognosis and increased metastatic risk^20^, suggesting that immune activity might persist within this lesion despite spatial exclusion from malignant tissue, a feature that would be obscured by the spatial topology analysis at the cell type level alone.

**Figure 5.**
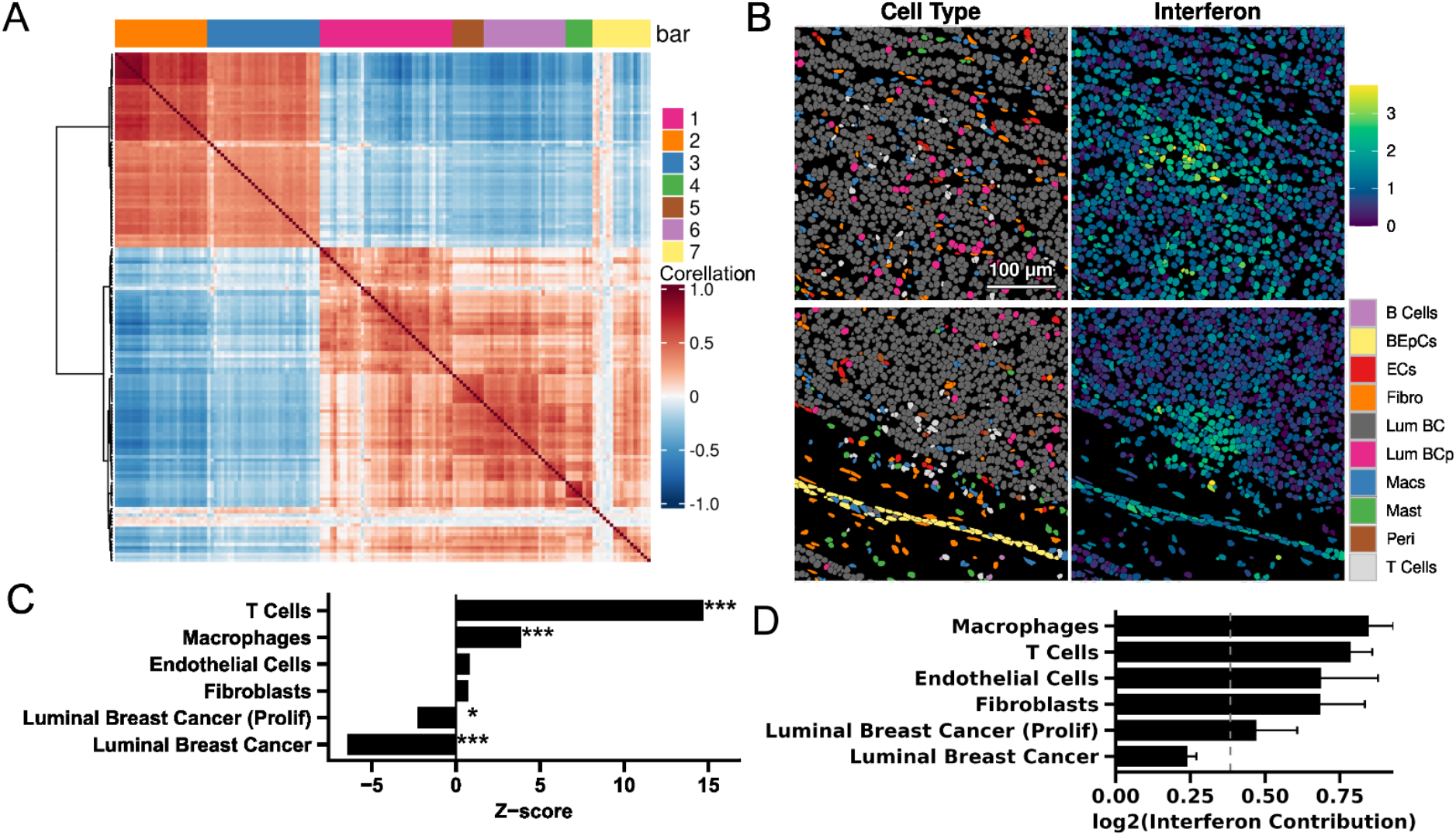
Spatial gene co-expression analysis. **A**. Heatmap depicting spatial gene co-expression modules **B**. Spatial plot showing metagene expression from cluster 7 (right) and equivalent cell types (left). **C**. Bar plot showing enrichment of cell type frequency within interferon hubs compared to frequency in malignant tissue regions as a whole. Positive Z-scores indicates increased frequency within interferon regions and negative Z-scores represents decreased frequency. * p < 0.05, ** p < 0.01, *** p < 0.001 (Benjamini-Hochberg corrected) **D**. Bar chart showing the average contribution of cell types to the interferon response gene signature. Error bars represent standard deviation of interferon metacluster expression in all cells for the relevant cell type.

## Discussion

Spatial transcriptomics holds immense potential for advancing precision oncology, yet barriers to tissue acquisition have limited its clinical translation. Here, we demonstrated that routinely discarded tissue sections from clinical breast cancer biopsies yield high-quality spatial transcriptomic data that recapitulate clinical diagnoses while providing enhanced characterization of the tumor microenvironment. From a single discarded section analyzed with a 493-gene panel, we identified over 50,000 cells, characterized diverse immune and stromal populations, and uncovered spatial features relevant to breast cancer biology including immune exclusion patterns and localized interferon response signatures. Critically, this approach circumvents major barriers to tissue access by repurposing material that would otherwise be discarded. This eliminates the need for patients to undergo additional invasive procedures exclusively for research, while seamlessly integrating into existing clinical workflows. This framework could enable an easier and more systematic collection of spatial data across diverse patient populations, addressing a key bottleneck in building the large-scale datasets required for developing spatially-informed diagnostic and prognostic tools.

Advancing precision oncology requires large-scale patient datasets, yet acquiring tissue for research remains a major bottleneck. Patient consent for additional biopsies is hindered by multiple factors including distrust in tissue use, particularly among marginalized groups^21,22^, reluctance to undergo additional invasive procedures^23,24^, and logistical barriers in clinical workflows. Another key barrier is the availability of treatment-naïve tissue samples, where CNBs are typically the sole source of such tissue samples. Patients with triple negative breast cancer and most patients with HER2-positive breast cancer (particularly those with stage II-III disease) often receive neoadjuvant treatment immediately post-diagnosis^25^. Our approach fundamentally addresses these barriers by utilizing treatment-naïve tissue that is already collected for clinical purposes and would otherwise be discarded. Because no additional tissue collection is required, this framework eliminates concerns about invasive procedures and reduces the consent burden on patients and overcomes situations where patients consent to research biopsies but have insufficient tissue for additional collections. Additionally, by performing spatial transcriptomics on the same biopsy used for diagnosis, we remove heterogeneity bias between clinical annotations and research findings. Moreover, the predefined analytical workflow and direct clinical relevance of spatial transcriptomic data enable clear communication about tissue use, potentially increasing transparency and trust. This approach is particularly advantageous for building diverse, representative biobanks, as it lowers patient-dependent barriers that disproportionately affect marginalized group participation^26^. By maximizing the scientific value of tissue already obtained through standard clinical care, this framework could dramatically expand access to primary samples while respecting patient autonomy and addressing historical inequities in biomedical research.

Clinical translation of novel technologies faces substantial barriers, with the average time from scientific discovery to routine clinical implementation exceeding 17 years^27^. A primary obstacle is workflow disruption— new methods that require substantial changes to established protocols face resistance and slow adoption. Our approach addresses this challenge by integrating seamlessly into existing histopathological workflows. The only modification required is transferring discarded tissue sections onto specialized slides rather than disposal, a step that adds one additional slide collection to routine processing and requires no specialized training beyond standard slide handling. Importantly, we achieved high-quality spatial transcriptomic data without implementing additional RNAse-free precautions beyond standard clinical histology protocols, demonstrating that routine clinical processing is sufficient to preserve RNA integrity for spatial analysis. From this single section, spatial transcriptomics provides subcellular-resolution characterization of the tumor microenvironment, including cell type identification and transcriptomic profiling across thousands of cells. The minimal workflow modifications required, combined with the substantial information gained, position this approach as a practical pathway to understand how advanced molecular profiling can be integrated into routine diagnostic pathology.

The development of clinically validated prognostic tests exemplifies the value of large-scale patient datasets for precision oncology. For instance, gene expression assays such as OncotypeDX, Mammaprint, and PAM50 have become widely adopted in breast cancer management, established through multi-institutional studies that identified expression patterns associated with clinical outcomes and treatment response^28–30^. However, such tests rely on bulk tissue analysis and do not capture the spatial architecture of the tumor microenvironment, which increasingly appears critical for predicting patient outcomes across cancer types^31^. Current diagnostic approaches require multiple tissue sections with individual stains for each marker^32^, an approach that is both resource-intensive and unable to capture the complex, spatially-organized interactions within tumors. Developing the next generation of spatially-informed prognostic tools, analogous to these gene expression-based approaches but incorporating spatial context, requires systematic collection of spatial transcriptomic data linked to longitudinal clinical outcomes across diverse patient populations. Our framework is uniquely positioned to enable such efforts by providing high-quality spatial data from treatment-naïve clinical biopsies that are directly tied to pathological diagnoses. By repurposing discarded tissue from routine diagnostic procedures, this approach could facilitate the rapid accrual of large, clinically-annotated spatial datasets necessary for discovering novel biomarkers, validating spatial prognostic signatures, and ultimately advancing precision medicine.

The integration of AI and traditional machine learning into diagnostic pathology represents a major opportunity in precision oncology^33,34^. Current AI approaches predominantly leverage morphological features from H&E-stained images to identify tissue patterns^35^, which can augment pathologist annotations but provide limited insight into the underlying molecular biology of the tumor microenvironment^36^. While morphology-based methods can identify broad immune cell populations^37^, they cannot distinguish functional cell subtypes or characterize their transcriptional states—information that offers the potential for predicting treatment response and patient outcomes that requires multiple cell-type-specific immunostains and is impractical to obtain comprehensively^38^. Spatial transcriptomics fundamentally addresses this limitation by providing spatially-resolved molecular profiles alongside tissue architecture in a single assay. For instance, our analysis revealed discrete spatial hubs of interferon response signatures involving coordinated expression of chemokines, interferon-stimulated genes, and immune regulators—activity associated with antitumor immunity^19,20^, that would be invisible to morphological analysis alone and cannot be captured by any single immunostain. AI models trained on such multimodal data could identify prognostic signatures, predict therapeutic responses and stratify patients for treatment in ways that morphology-based approaches cannot achieve, therefore augmenting current diagnostic tools such as OncotypeDX. By integrating spatial transcriptomic data with traditional histopathology from the same biopsy, this framework provides a scalable approach for systematically collecting the clinically-annotated datasets necessary for training and validating these next-generation AI diagnostic tools across diverse patient populations. Future implementation of agentic spatial data analysis will not only allow for the semi-automatic generation of diagnostic reports but could also immediately connect patients to ongoing clinical trials and cutting-edge research. Altogether, it represents a critical step toward achieving the goals of true precision medicine

## Conclusion

Spatial transcriptomics holds transformative potential for precision oncology and here we demonstrate that routinely discarded sections from clinical biopsies yield high-quality spatial transcriptomic data. These results reveal cellular composition, spatial architecture, and molecular features—such as immune exclusion patterns and interferon response hubs—that are relevant to cancer biology and clinical outcomes. Critically, this approach can integrate seamlessly into existing diagnostic workflows without requiring additional tissue from patients, thereby addressing major barriers related to consent, tissue availability, and clinical feasibility. By maximizing the scientific value of material that would otherwise be discarded, this framework could enable the systematic collection of spatially-resolved, clinically-annotated data across diverse patient populations. Such datasets are essential for developing the next generation of spatially-informed diagnostic tools, validating prognostic biomarkers, and ultimately realizing the promise of truly personalized cancer medicine.

## Methods

### Patient sample collection

This study was conducted with approval from the Boston University (BU) Institutional Review Board (IRB), ensuring adherence to ethical standards. Informed consent was obtained from each participant prior to enrollment. All ethical regulations relevant to human research participants were followed (H-38024 and H-38678).

### Biopsy processing

The biopsy was collected from the patient and within 5 minutes was placed in 10% neutral buffered formalin for 7 hours and 40 minutes. The sample was processed according to standard histopathology workflow. A single 5 µm thick section was placed onto a functionalized Spatial Genomics slide within the imageable region then dried at 58°C for 1 hour (**Supplemental Fig. 2B**) and stored at −20°C.

### Immunohistochemistry

Immunohistochemistry was performed on the formalin-fixed, paraffin-embedded (FFPE) sections, stained for ER, PgR, and Ki-67 on a clinical autostainer according to ASCO/CAP guidelines against receptor clones: SP1, 1E2, and 30-9, respectively. The stained slides were digitized on the Leica Aperio AT2 at 40x magnification.

### seqFish workflow

*In situ* spatial transcriptomics was performed using the Spatial Genomics Gene Positioning System (GenePS). The samples were hybridized with the Spatial Genomics Human Immuno-Oncological Panel. The automated GenePS platform was used for image acquisition, reagent delivery, and data processing. The instrument was set up following the *GenePS Instrument User Guide (INS-205)*. Regions of interest were identified from 10x images. Post-imaging and ROI selection, the automated imaging protocol imaged the sample with a 60x oil objective, capturing six z-planes at 1.5 µm z-steps. The slide underwent iterative cycles of decode probe hybridization, imaging and signal removal. Raw images were aligned across the different rounds of hybridization and processing was performed on the GenePS instrument to identify RNA signals. Transcripts were decoded using the Spatial Genomics Navigator software and a maximum false discovery rate of 5% was applied.

### Cell segmentation

DAPI images were scaled to pixel sizes of 0.5 μm. Individual nuclei were identified from this scaled image using cellpose^39^. A gene map consisting of all genes was created by adapting methods described by the UCS method for the transcript coordinates outputted by the GenePS platform^40^. Cell segmentation was performed using UCS on the nuclei segmentation and the gene map as described^40^.

### H&E nuclei segmentation

Clinical H&E slides were digitized on the Leica Aperio AT2 at 40x magnification. Nuclei were segmented from whole slide images for the clinically H&E-stained images using QuPath (v0.5.1) with default parameters in the ‘*Cell Detection*’ module.

### Pre-processing seqFISH data

All spatial omics data processing was performed using the Giotto Suite platform unless specified otherwise (v4.2.1) in R (v4.4.0)^41,42^. Cells were defined based on the segmentation performed by UCS and imported into Giotto using ‘*createGiottoPolygonsFromMask*’ then scaled back to the extents using ‘*rescale*’. Transcripts were used as exported by the GenePS platform and imported into Giotto using ‘*createGiottoPoints*’. Cell polygons and transcripts were integrated into a Giotto object using ‘*createGiottoObjectSubcellular*’ and transcripts were assigned to cells through overlap to cell polygons with ‘*calculateOverlapRaster*’, *overlapToMatrix*’, and ‘‘addSpatialCentroidLocations’. Cells were filtered with ‘*filterGiotto’* to remove cells that contained less than 25 unique transcripts. Transcripts were removed if they were found in less than 100 cells and were normalized using the default parameters with ‘*normalizeGiotto*’.

### Clustering

PCA was calculated based on all transcripts with ‘*runPCA*’ using default parameters and the UMAP representation was calculated using the top 13 PCs using ‘*runUMAP*’. A nearest neighbor network was calculated with ‘*createNearestNetwork*’ using the top 13 PCs and a k of 13. Leiden clustering was performed with ‘*doLeidenCluster*’ using default parameters with a resolution of 0.2 and 1000 iterations. From the 9 clusters identified scran with ‘*findMarkers_one_vs_all*’ with default parameters was used to determine marker genes based on the transcript rank, then cell types were determined manually. Clusters 1 and 2 were merged to both be “Luminal epithelial cancer cells” based on gene expression profiles.

### Marker gene visualization

Cells were separated into cancer (luminal breast cancer cells and proliferative luminal breast cancer cells) and non-cancer. To account for issues with cell segmentation not encompassing all transcripts, marker transcripts were associated with the closest cell, within a one cell diameter. Cells were considered positive for a marker if expression was greater than 1 transcript per cell for the relevant marker. Spatial maps of cells were created with ‘*spatInSituPlotPoints*’ and cells were colored based on a binary for being positive or negative for a marker then visually compared to stains conducted for patient diagnosis. Proportions were calculated based on the total number of cells expressing each marker within the cancer population as determined through immunohistochemistry (IHC) in accordance with standard operating procedures at Boston Medical Center. The pathology expression percentages were taken from the patient’s pathology report from diagnosis. Positivity for ESR1 and PGR were set at greater than 10% and less than 17% was considered low for Ki67.

### Cell-cell interaction analysis

The cell-cell interaction was performed first by creating a Delaunay spatial network with ‘*createSpatialNetwork*’ using default parameters with a minimum k of 2 and maximum distance of 20 µm. Cell proximity enrichment was calculated with ‘*cellProximityEnrichment*’ using default parameters to identify cell-cell interactions. Niche clustering was used to identify spatially organized patterns of cells using ‘*calculateSpatCellMetadataProportions’* with a Delaunay network followed by *‘getSpatialEnrichment’* with default parameters to calculate enrichment. Specific niches were identified using *‘kmeans’* with 6 centers and 100 max iterations. Proportions for niche composition were calculated based on the total number of cells in the respective niche.

### Gene co-expression analysis

Spatially variable genes were identified using ‘*binSpect*’ with a kmeans binning method and Delaunay network. The top spatial genes were selected based on having a corrected p value of less than 0.001 and a score greater than 1000. Using the top spatial genes, spatially correlated transcripts were identified using ‘*detectSpatialCorFeats*’ using a kNN nearest neighbor network. The spatial correlations were clustered using ‘*clusterSpatialCorFeats*’ with a k of 7. Annotations of genes cluster modules were performed manually based on the transcripts found in each module of spatially correlated features. Expression of each module was calculated per cell using ‘*createMetafeats*’ to create metafeatures that represented all transcripts within the module. These expression values were used to visualize the spatial distribution of each spatial module. The contribution of each individual cell type was calculated as a mean of expression within each cell type for the relevant module.

### Identification of interferon-responsive microenvironments within cancer regions

Cancer-enriched tissue regions were defined by constructing a Delaunay spatial network with ‘createSpatialNetwork’ restricted to cancer cells with a maximum distance of 20 µm. Subnetworks were extracted using a minimum size of 2000 cells. Concave hull boundaries were computed around these subnetworks to delineate cancer portions. Cells whose polygons intersected these boundaries were assigned to their respective cancer portions. Within cancer portions, interferon hotspots were identified by selecting cells with high interferon signaling (cluster metagene 7 ≥ 1.75) and extracting their spatial neighbors using a Delaunay network with a maximum distance of 10 µm to ensure proximity. Contiguous interferon hotspots were extracted using a minimum size of 25 cells that all meet the “high interferon signaling” criteria. To test for cell type enrichment in interferon hotspots, observed cellular composition was compared to a null distribution generated by 1000 permutations of randomly sampling spatially connected, cancer-dense regions of equivalent size from within cancer portions. Enrichment was quantified using z-scores, with significance assessed after Benjamini-Hochberg correction (α = 0.05).

## Acknowledgments

We are grateful to the patient who generously donated their sample for this study. We also thank Dr. Christopher D. Andry (Department of Pathology and Laboratory Medicine, Boston Medical Center, Boston University Chobanian & Avedisian School of Medicine) for his valuable advice throughout this project.

## Author Contributions

J.P.S., J.G.C., N.Y.K., and R.D. conceived and designed the study. J.P.S., V.J., A.J., and I.A. performed the analysis and data curation. L.S.^1^ and T.N. contributed to the wet lab work and methodology. J.P.S. drafted the original manuscript. Q.S., Y.W., J.G.C., S.B.M., E.J.B., L.S.^2^, C.M.H., N.Y.K., and R.D. contributed to manuscript review and editing. N.Y.K. and R.D. provided supervision, project administration, and funding acquisition. All authors reviewed and approved the final manuscript.

## References

1. Zhou, Y. et al. Tumor biomarkers for diagnosis, prognosis and targeted therapy. Signal Transduct. Target. Ther. 9, 132 (2024).

2. Zou, J. & Wang, E. Cancer Biomarker Discovery for Precision Medicine: New Progress. Curr. Med. Chem. 26, 7655–7671.

3. Sakai, S. A. et al. Single-cell spatial analysis with Xenium reveals anti-tumour responses of CXCL13 + T and CXCL9+ cells after radiotherapy combined with anti-PD-L1 therapy. Br. J. Cancer 133, 795–808 (2025).

4. Klughammer, J. et al. A multi-modal single-cell and spatial expression map of metastatic breast cancer biopsies across clinicopathological features. Nat. Med. 30, 3236–3249 (2024).

5. Janesick, A. et al. High resolution mapping of the tumor microenvironment using integrated single-cell, spatial and in situ analysis. Nat. Commun. 14, 8353 (2023).

6. Schott, M. et al. Open-ST: High-resolution spatial transcriptomics in 3D. Cell 187, 3953–3972.e26 (2024).

7. Pentimalli, T. M., Karaiskos, N. & Rajewsky, N. Challenges and Opportunities in the Clinical Translation of High-Resolution Spatial Transcriptomics. Annu. Rev. Pathol. Mech. Dis. 20, 405–432 (2025).

8. Bar, Y. et al. Selection biases in the systematic collection of breast biobank specimens. NPJ Breast Cancer 11, 83 (2025).

9. Koh, S.-B. et al. Systematic tissue collection during clinical breast biopsy is feasible, safe and enables high-content translational analyses. Npj Precis. Oncol. 5, 85 (2021).

10. Allott, E. H. et al. Intratumoral heterogeneity as a source of discordance in breast cancer biomarker classification. Breast Cancer Res. 18, 68 (2016).

11. Harati, M. D. et al. An introduction to starting a biobank. Methods Mol. Biol. Clifton NJ 1897, 7–16 (2019).

12. Eng, C.-H. L. et al. Transcriptome-scale super-resolved imaging in tissues by RNA seqFISH+. Nature 568, 235–239 (2019).

13. Zhang, S. Y. et al. E2F-1: A Proliferative Marker of Breast Neoplasia1. Cancer Epidemiol. Biomarkers Prev. 9, 395–401 (2000).

14. Liang, X., Li, L. & Fan, Y. Diagnostic, Prognostic, and Immunological Roles of HELLS in Pan-Cancer: A Bioinformatics Analysis. Front. Immunol. 13, 870726 (2022).

15. Grünwald, B. T. et al. Spatially confined sub-tumor microenvironments in pancreatic cancer. Cell 184, 5577–5592.e18 (2021).

16. Meylan, M. et al. Tertiary lymphoid structures generate and propagate anti-tumor antibody-producing plasma cells in renal cell cancer. Immunity 55, 527–541.e5 (2022).

17. Wang, X. et al. Spatial transcriptomics reveals substantial heterogeneity in triple-negative breast cancer with potential clinical implications. Nat. Commun. 15, 10232 (2024).

18. Wu, B., Zhang, B., Li, B., Wu, H. & Jiang, M. Cold and hot tumors: from molecular mechanisms to targeted therapy. Signal Transduct. Target. Ther. 9, 274 (2024).

19. Andersson, A. et al. Spatial deconvolution of HER2-positive breast cancer delineates tumor-associated cell type interactions. Nat. Commun. 12, 6012 (2021).

20. Lamsal, A. et al. Opposite and dynamic regulation of the interferon response in metastatic and non-metastatic breast cancer. Cell Commun. Signal. 21, 50 (2023).

21. Dang, J. H. T. et al. Engaging diverse populations about biospecimen donation for cancer research. J. Community Genet. 5, 313–327 (2014).

22. Drake, B. F., Boyd, D., Carter, K., Gehlert, S. & Thompson, V. S. Barriers and strategies to participation in tissue research among African-American men. J. Cancer Educ. Off. J. Am. Assoc. Cancer Educ. 32, 51–58 (2017).

23. Braun, K. L. et al. Cancer Patient Perceptions about Biobanking and Preferred Timing of Consent. Biopreservation Biobanking 12, 106–112 (2014).

24. Lewis, C. et al. Public views on the donation and use of human biological samples in biomedical research: a mixed methods study. BMJ Open 3, e003056 (2013).

25. Gradishar, W. J. et al. Breast Cancer, Version 3.2024, NCCN Clinical Practice Guidelines in Oncology. J. Natl. Compr. Canc. Netw. 22, 331–357 (2024).

26. Dang, J. H. T. et al. Engaging diverse populations about biospecimen donation for cancer research. J. Community Genet. 5, 313–327 (2014).

27. Morris, Z. S., Wooding, S. & Grant, J. The answer is 17 years, what is the question: understanding time lags in translational research. J. R. Soc. Med. 104, 510–520 (2011).

28. Siow, Z. R., De Boer, R. H., Lindeman, G. J. & Mann, G. B. Spotlight on the utility of the Oncotype DX® breast cancer assay. Int. J. Womens Health 10, 89–100 (2018).

29. Vijver, M. J. van de et al. A Gene-Expression Signature as a Predictor of Survival in Breast Cancer. N. Engl. J. Med. 347, 1999–2009 (2002).

30. Parker, J. S. et al. Supervised Risk Predictor of Breast Cancer Based on Intrinsic Subtypes. J. Clin. Oncol. 27, 1160–1167 (2009).

31. Arora, R. et al. Spatial transcriptomics reveals distinct and conserved tumor core and edge architectures that predict survival and targeted therapy response. Nat. Commun. 14, 5029 (2023).

32. Gurina, T. S. & Simms, L. Histology, Staining. in StatPearls (StatPearls Publishing, Treasure Island (FL), 2025).

33. Ma, Y., Jamdade, S., Konduri, L. & Sailem, H. AI in Histopathology Explorer for comprehensive analysis of the evolving AI landscape in histopathology. Npj Digit. Med. 8, 156 (2025).

34. Försch, S., Klauschen, F., Hufnagl, P. & Roth, W. Artificial Intelligence in Pathology. Dtsch. Ärztebl. Int. 118, 199–204 (2021).

35. Komura, D. & Ishikawa, S. Machine Learning Methods for Histopathological Image Analysis. Comput. Struct. Biotechnol. J. 16, 34–42 (2018).

36. Greeley, C., Holder, L., Nilsson, E. E. & Skinner, M. K. Scalable deep learning artificial intelligence histopathology slide analysis and validation. Sci. Rep. 14, 26748 (2024).

37. Nakkireddy, S. R. et al. Integrative analysis of H&E and IHC identifies prognostic immune subtypes in HPV related oropharyngeal cancer. Commun. Med. 4, 190 (2024).

38. Chiang, C.-L. et al. Clinical utility of immunohistochemical subtyping in patients with small cell lung cancer. Lung Cancer 188, 107473 (2024).

39. Stringer, C., Wang, T., Michaelos, M. & Pachitariu, M. Cellpose: a generalist algorithm for cellular segmentation. Nat. Methods 18, 100–106 (2021).

40. Chen, Y., Xu, X., Wan, X., Xiao, J. & Yang, C. UCS: A Unified Approach to Cell Segmentation for Subcellular Spatial Transcriptomics. Small Methods 9, 2400975 (2025).

41. Chen, J. G. et al. Giotto Suite: a multiscale and technology-agnostic spatial multiomics analysis ecosystem. Nat. Methods 1–13 (2025) doi:10.1038/s41592-025-02817-w.

42. R Core Team. R: A Language and Environment for Statistical Computing. R Foundation for Statistical Computing (2020).

