## supplemental figures for "Clinical feasibility of spatial transcriptomics using discarded tissue from diagnostic breast biopsies"

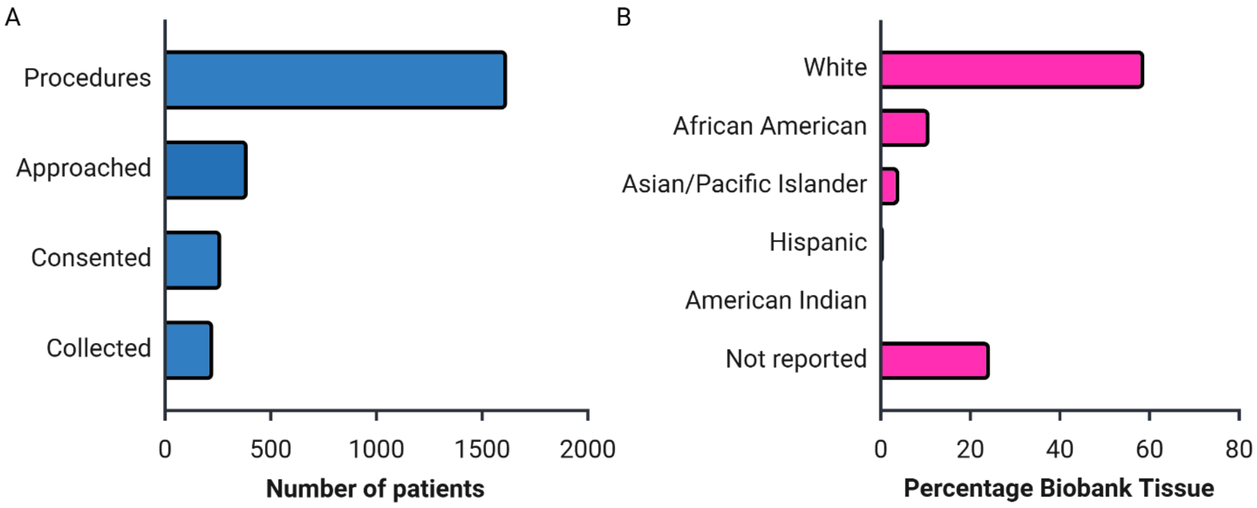


**Supplemental Figure 1. Biobanking collection statistics and limitations**. **A.** Bar chart demonstrating the success of subsequent tissue collection steps for all eligible patients undergoing procedures, data obtained from Koh *et al* 2021. **B.** Ethnicity distribution for all biobanked tissue in Bar *et al* 2025.

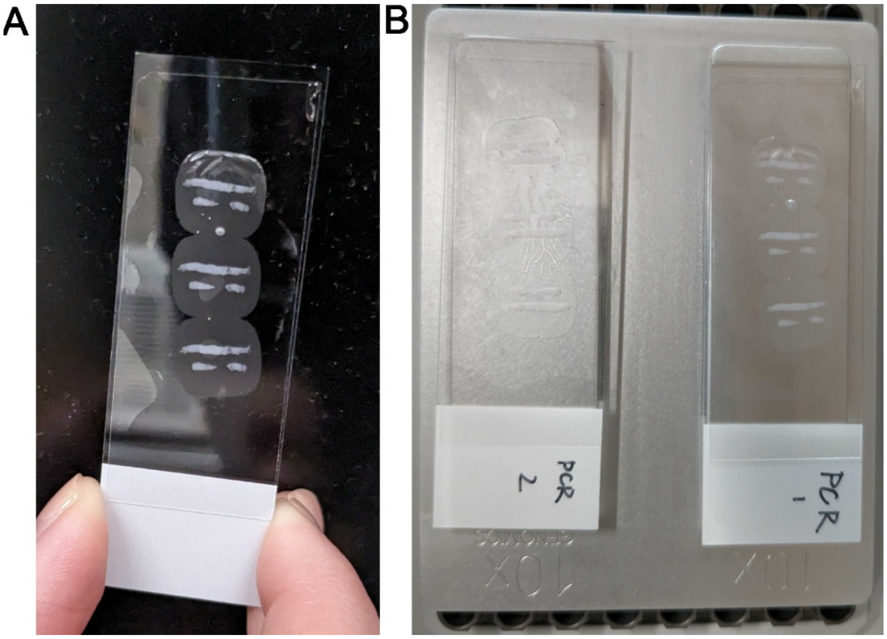


**Supplemental Figure 2. Tissue placement on specialized seqFISH slides. A**. Image from the seqFISH (Spatial Genomics) glass slide used for collecting the discarded tissue sections. **B.** Image of obtained slides after heating at 58°C for 1 hour prior to storage at -20°C until seqFISH was performed

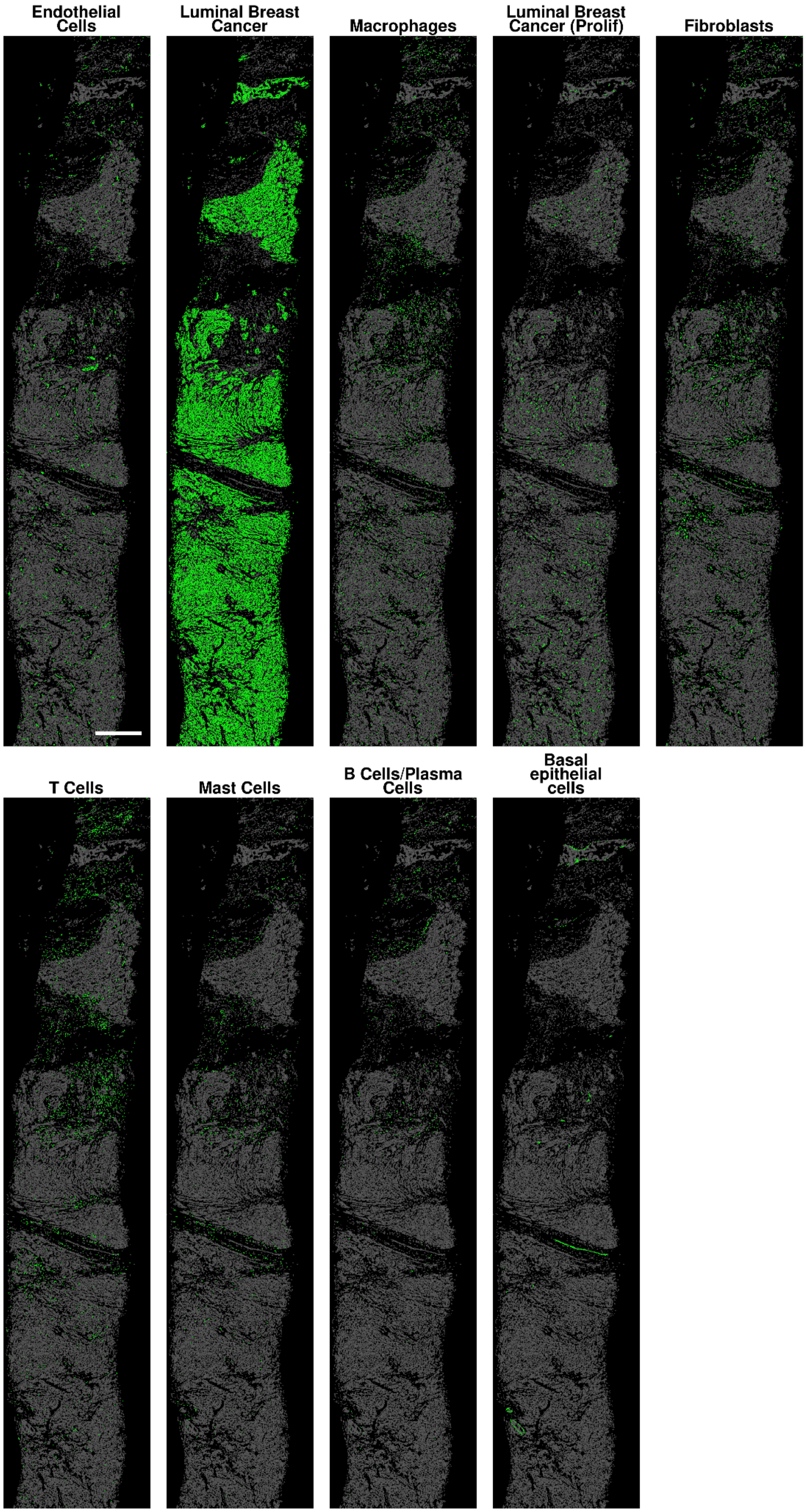


**Supplemental Figure 3**. **Individual cell type spatial maps**. Spatial plots of biopsy tissue (green = indicated cell type, grey = all other cell types). Scale bar represents 500 μm.

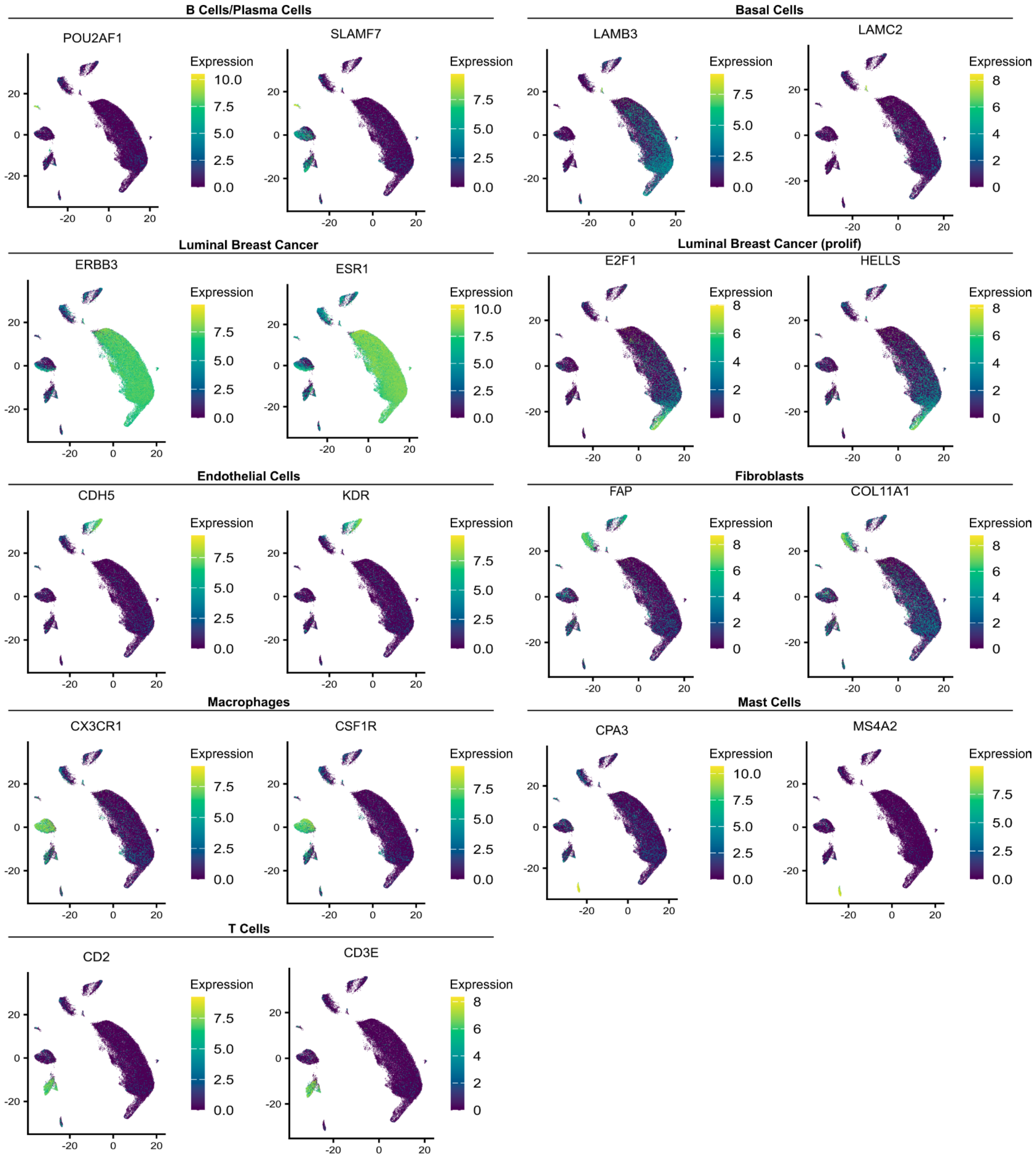


**Supplemental Figure 4**. **Marker gene expression across all cells in UMAP space.** Uniform Manifold Approximation and Projection (UMAP) of all cells showing marker gene expression for each cell type. Individual points are cells colored by expression levels of the relevant marker. Blue and yellow indicate low and high expression, respectively.

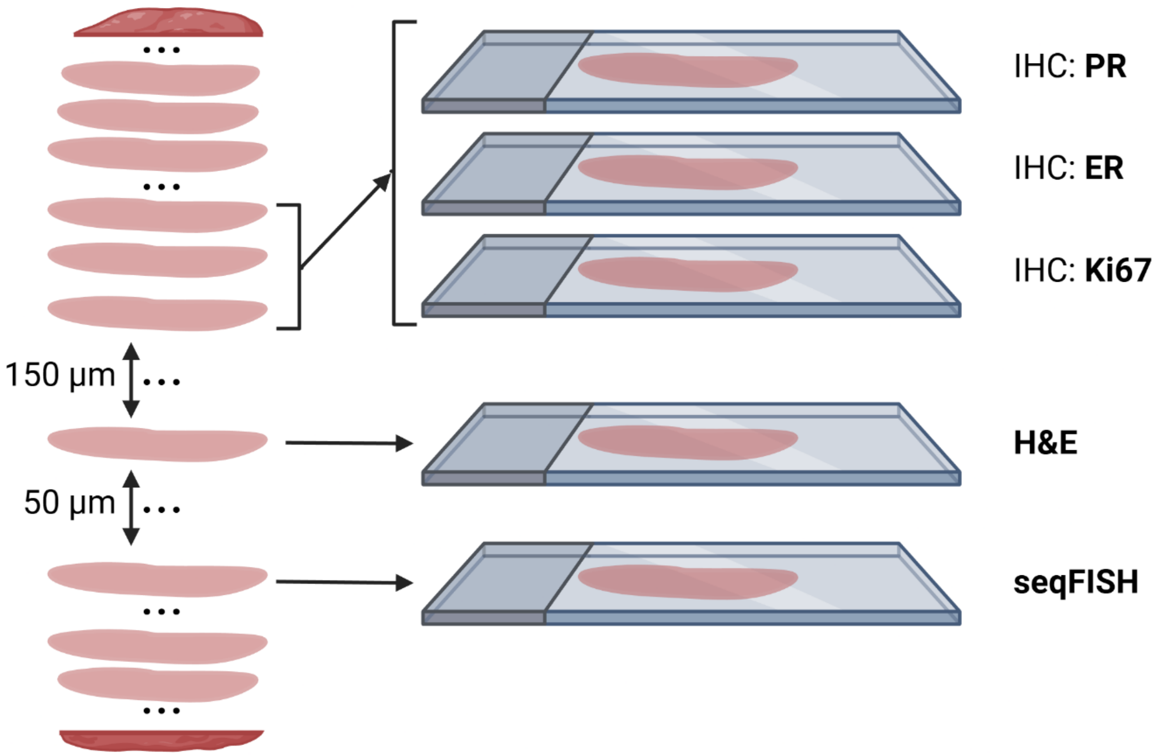


**Supplemental Figure 5. Tissue collection process from the diagnostic biopsy**. Schematic representation demonstrating how tissue sections were obtained for pathology (PR, ER, Ki67, and H&E) and research (seqFISH). Not shown are the multiple levels of collection typically performed during biopsy processing.

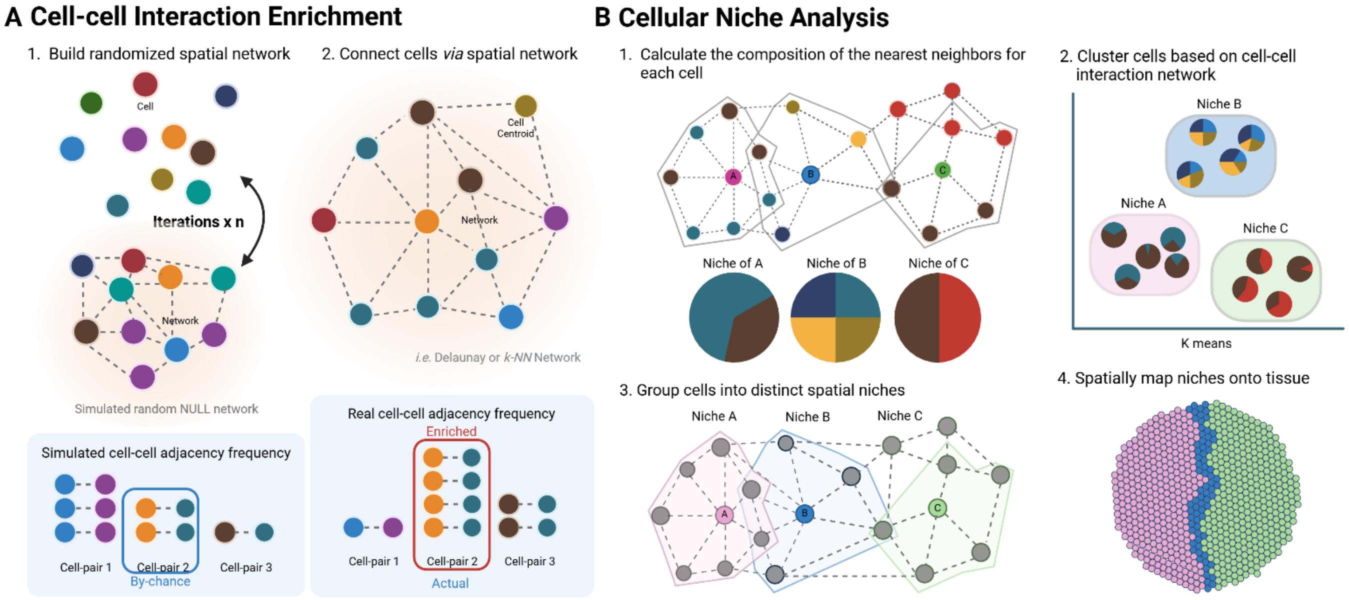


**Supplemental Figure 6**. **Schematic overview of cell-cell interactions methods**. Cell-cell interaction enrichment (**A**) and cellular niche analysis (**B**)

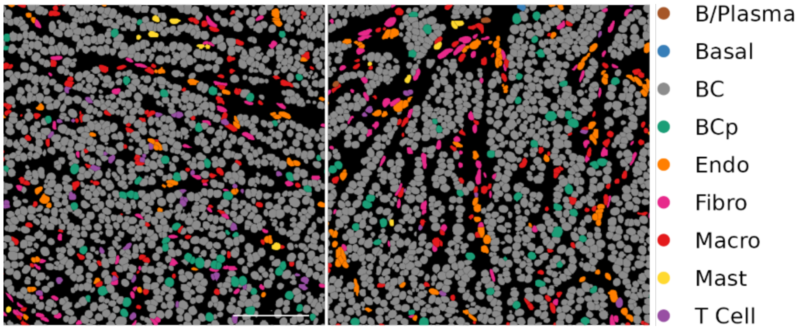


**Supplemental Figure 7**. **Spatial distribution of immune cells**. Spatial map showing different tissue regions within the biopsy and cells are colored by cell type. B Cell/Plasma Cell (B/Plasma), basal cells (Basal), luminal breast cancer (BC), proliferative luminal breast cancer (BCp), endothelial (Endo), fibroblasts (Fibro), macrophages (Macro), mast cells (Mast). Scale bar represents 100 μm.

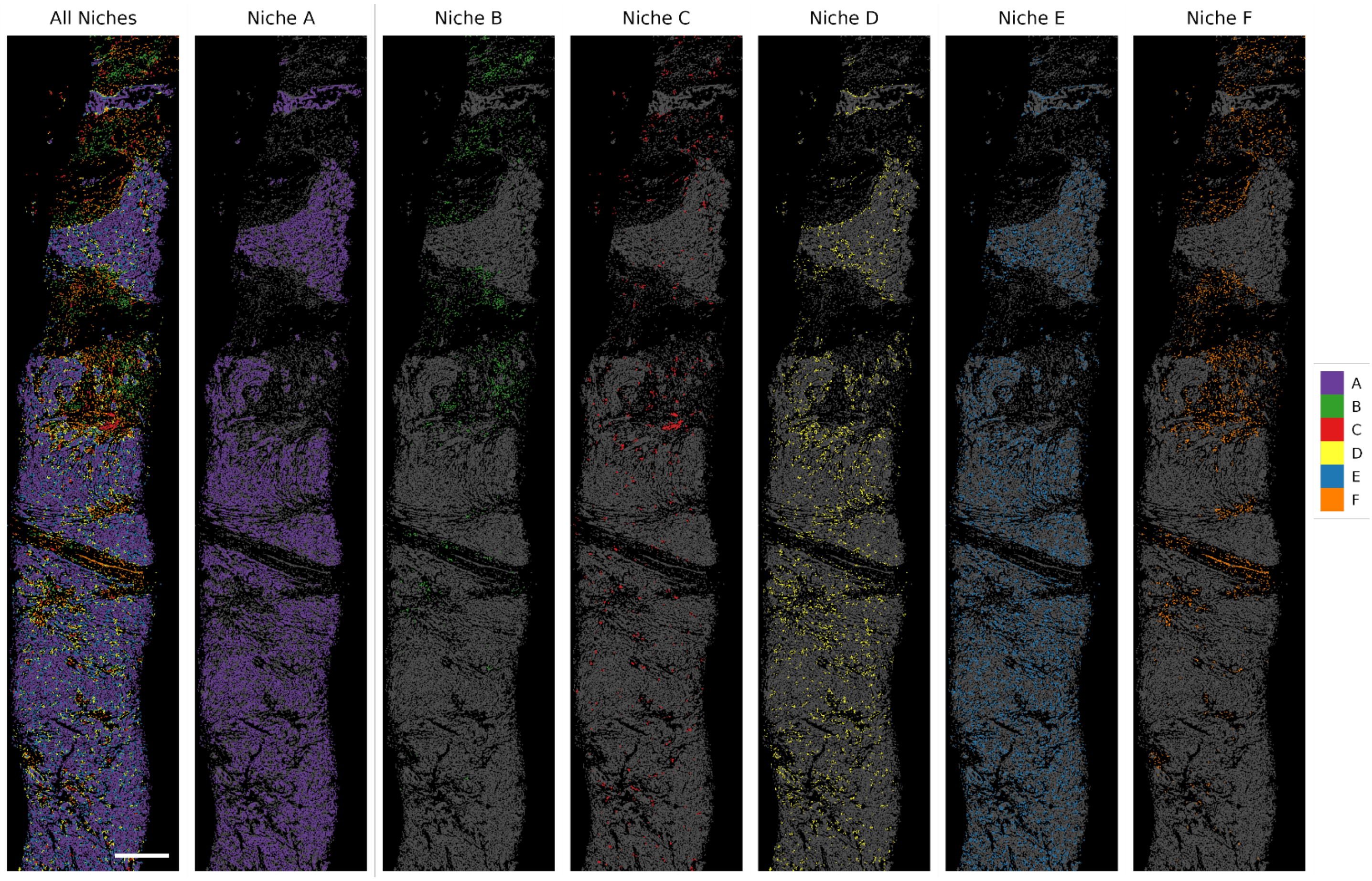


**Supplemental Figure 8. Spatial map of cellular niches**. Panel on the left shows all identified cellular niches within the biopsy. Subsequent panels show each individual niche colored by its assigned color, while other cells are colored grey. Scale bar represents 500 µm.

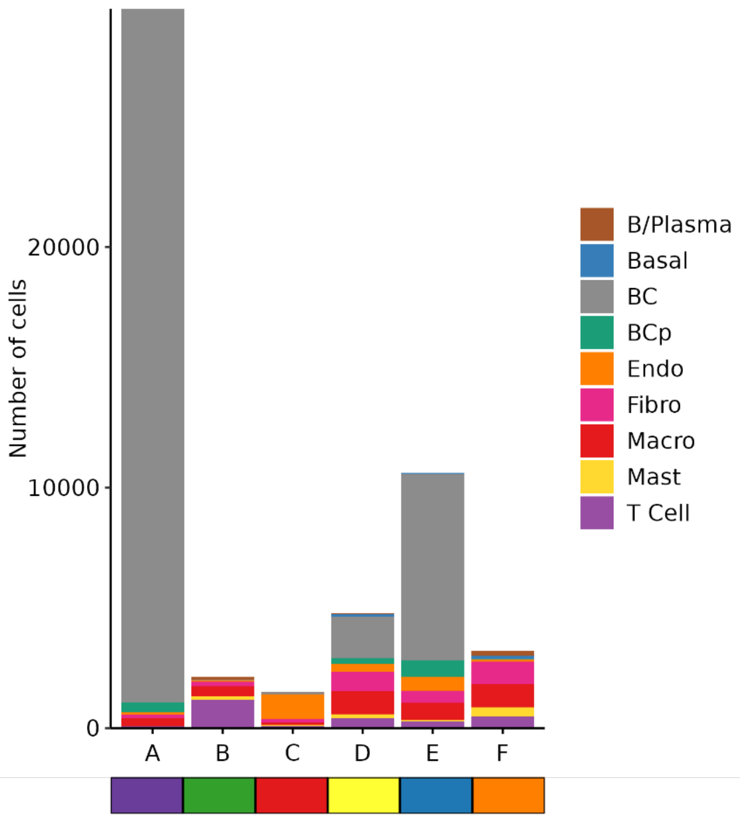


**Supplemental Figure 9. Raw cell composition of each spatial niche**. Stacked barplot showing cell type proportions for each cellular niche. Cell/Plasma Cell (B/Plasma), basal cells (Basal), luminal breast cancer (BC), proliferative luminal breast cancer (BCp), endothelial (Endo), fibroblasts (Fibro), macrophages (Macro), mast cells (Mast).

### Spatial Co-expression Analysis

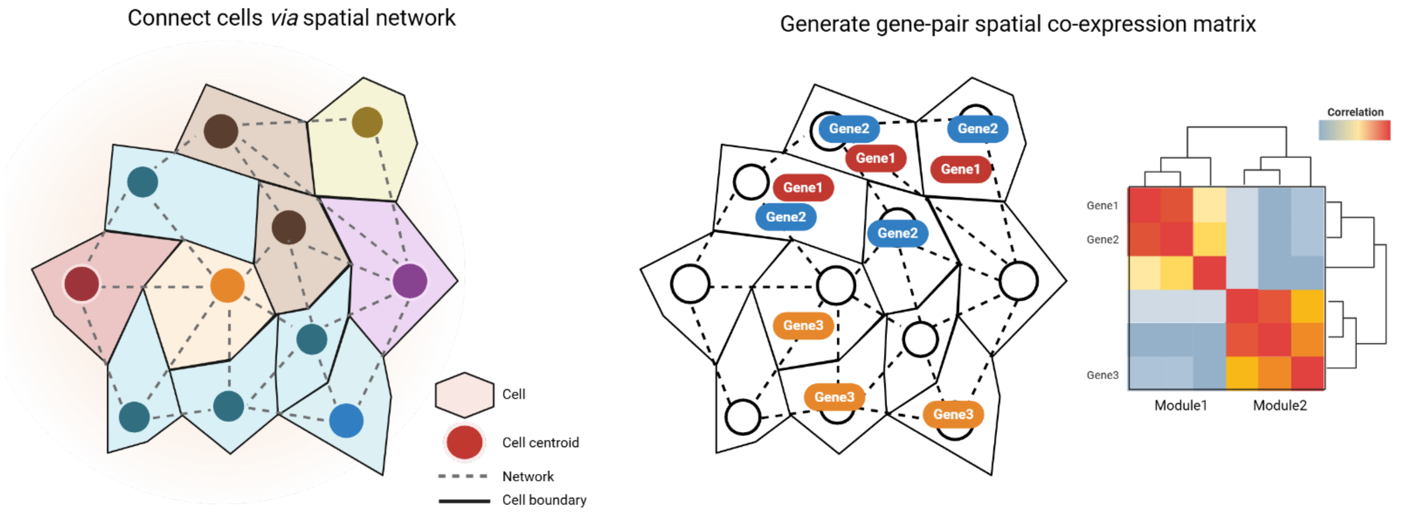


**Supplemental Figure 10**. **Schematic overview of the calculation of spatially co-expressed genes**. Left side represents the spatial network with each polygon and circle representing a single cell. Dotted lines indicate cell-cell interactions. Right side represents gene expression patterns with the spatial co-expression correlation indicating gene-pairs that are proximally expressed.

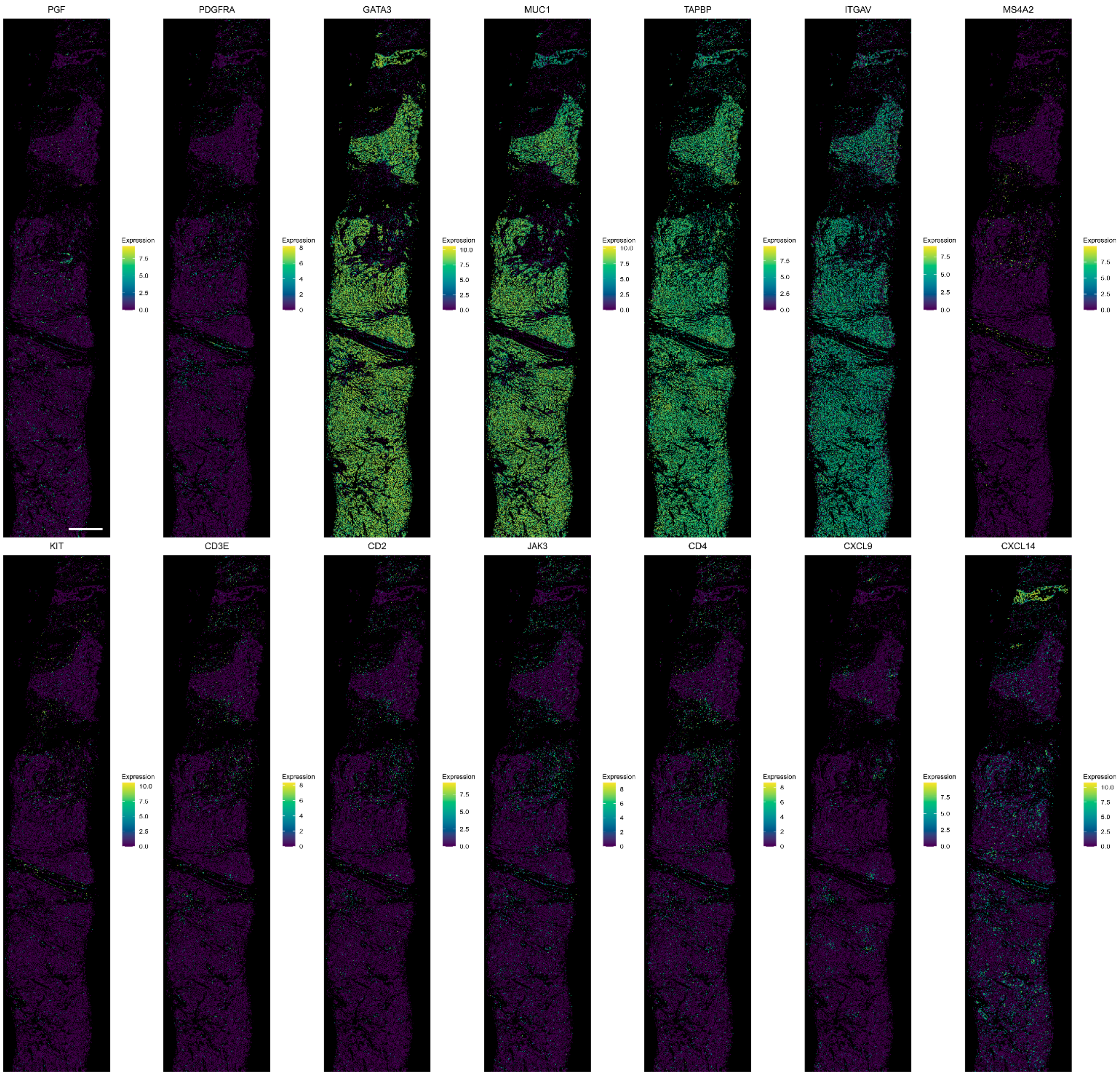


**Supplemental Figure 11. Spatial map of spatially organized genes.** Spatial plots showing the expression of the top 2 ranked spatially variable genes from each gene co-expression cluster. Blue and yellow indicate low and high expression, respectively.

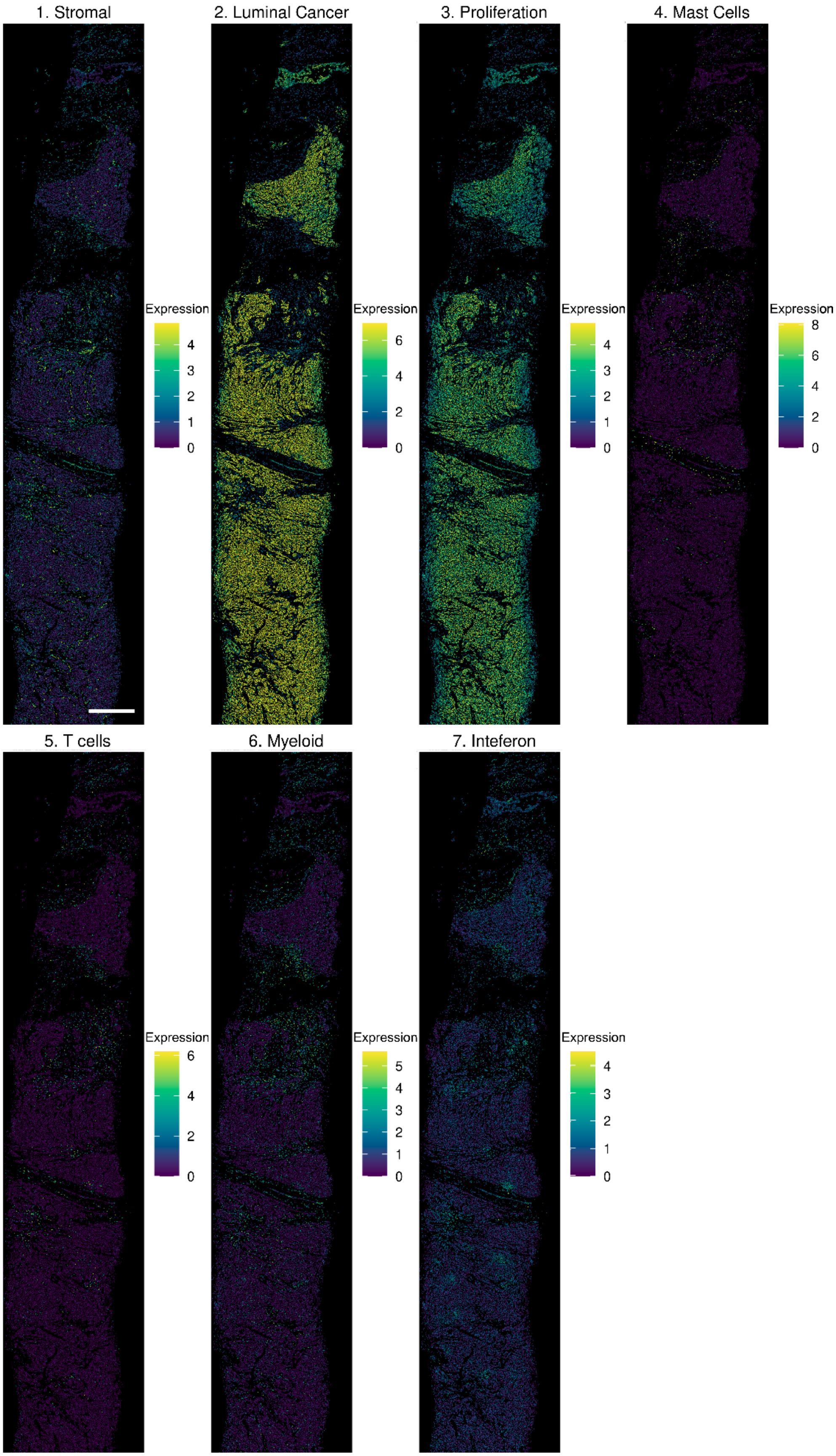


**Supplemental Figure 12. Spatial map of gene co-expression clusters**. Panels represent expression of gene metafeatures. Cells colored by expression of the aggregated metafeature. Blue and yellow indicate low and high expression, respectively.
